# Interactions between the microbiome and mating influence the female’s transcriptional profile in *Drosophila melanogaster*

**DOI:** 10.1101/2020.05.30.125427

**Authors:** Sofie Y. N. Delbare, Yasir H. Ahmed-Braimah, Mariana F. Wolfner, Andrew G. Clark

## Abstract

*Drosophila melanogaster* females undergo a variety of post-mating changes that influence their activity, feeding behavior, metabolism, egg production and gene expression. These changes are induced either by mating itself or by sperm or seminal fluid proteins. In addition, studies have shown that axenic females—those lacking a microbiome—have altered fecundity compared to females with a microbiome, and that the microbiome of the female’s mate can influence reproductive success. However, the extent to which post-mating changes in transcript abundance are affected by microbiome state is not well-characterized. Here we investigated fecundity and the post-mating transcript abundance profile of axenic or control females after mating with either axenic or control males. We observed interactions between the female’s microbiome and her mating status: transcripts of genes involved in reproduction and genes with neuronal functions were differentially abundant depending on the females’ microbiome status, but only in mated females. In addition, immunity genes showed varied responses to either the microbiome, mating, or a combination of those two factors. We further observed that the male’s microbiome status influences the fecundity of both control and axenic females, while only influencing the transcriptional profile of axenic females. Our results indicate that the microbiome plays a vital role in the post-mating switch of the female’s transcriptome.

## 1 Introduction

Reproductive success is determined by the cumulative effects of behavioral and physiological changes that a female undergoes after mating. In *Drosophila*, these post-mating responses include sperm storage, increased oocyte production and ovulation, a decrease in sleep and the female’s propensity to remate, alterations to the female’s immune system, and changes in feeding frequency, gut morphology and physiology (reviewed in [1]). These phenotypic changes are accompanied by extensive transcriptome changes across several female tissues [2–11]. These transcriptome changes typically reach their highest magnitude at around six hours after mating [5, 10], and often include genes that encode proteolytic/metabolic enzymes and immune response genes [5, 9–11]. Furthermore, these female post-mating responses are influenced by an interplay between the genotypes of the female and her mate [12–16], and are induced in part by male ejaculate components that are transferred to the female during mating [1, 4–6, 17, 18]. The post-mating changes in metabolism and food uptake are thought to be required to meet the high energetic demands of oocyte production [19, 20], and can be potentially be influenced by transient factors such as the host microbiome.

In recent years, *Drosophila melanogaster* has emerged as a valuable model to study fundamental principles of host-microbiome interactions, owing to the availability of genetic resources and a well-characterized and easily-manipulated gut microbiome [21]. Removing the microbiome (bacteria and yeast) from *D. melanogaster* affects a wide range of traits, including the gut transcriptome, which highlights the regulatory effects of the microbiome on tissue homeostasis, carbohydrate and lipid metabolism, proteolysis and immunity [22–26]. In addition, microbiome-induced transcriptome changes underlie a range of phenotypes such as larval development time [27–30], metabolite levels [30, 31], intestinal stem cell proliferation [32, 33], behavior [34, 35], longevity [28, 33, 36] and reproductive capacity [28, 37–39].

Across microbiome studies of *D. melanogaster*, some effects are consistently observed (e.g. transcriptome changes or changes in metabolite content), while others yield variable results, which likely depend on the experimental design and/or environmental conditions [21]. For example, Schretter *et al*. [35] observed a significant increase in the locomotor activity of axenic flies—those that lack a microbiome—relative to flies with a microbiome, but no significant difference in activity was seen by Selkrig *et al*. [40]. Furthermore, microbiome-induced changes in courtship were not observed by Selkrig *et al*. [40] and Leftwich et al. [41], while changes in courtship were observed by Qiao et al. [39] and Sharon *et al*. [42]. Even egg laying, which was consistently observed to be lower in axenic females compared to females with a microbiome in multiple studies [28, 37–40], was not observed to be lower in axenic females by Ridley *et al*. [30].

The varied results obtained in microbiome studies using *D. melanogaster* could be attributed to variability in nutrients [43–45], species and strains of microbiota and host, host age [46–48], or the requirement for frequent bacterial replenishment to maintain a stable microbiome [49, 50]. However, another variable that can play a role is female mating status. For example, several studies measured differences in the transcriptomes of female *D. melanogaster* with or without a microbiome, but if or when females mated was not explicitly controlled ([22], [23], [24], [26]). Thus, it is unclear how the interaction between microbiome and mating status influences whole-body transcript abundance in *D. melanogaster*.

Here we use short-read RNA sequencing (RNA-seq) to explore how the interaction between the female’s mating status (virgin or mated) and her microbiome state (control or axenic) influence her transcriptome. We further evaluate how the female transcriptome is influenced by the microbiome of the female’s mate, because previous work has shown that male mating success is impacted by his microbiome state [38]. Of particular interest is the post-mating up-regulation of immune response genes which occurs in females of a variety of species [51], including *D. melanogaster*. The post-mating up-regulation of some immunity genes was shown to be elicited by sperm and male seminal fluid proteins [4–6, 17], but it is unclear whether microbiome state influences post-mating immune gene up-regulation, even though activation of the innate immune system might seem particularly sensitive to prior microbial exposure.

## 2 Results

We investigated whether the female’s transcriptome is influenced by interactions between the female’s microbiome and her mating status. In addition, we asked whether the microbiome status of her mate influences her fecundity and post-mating transcriptional response. To this end, we mated wildtype Canton-S control females (containing a conventional microbiome found in our lab’s Canton-S stock) and axenic females (lacking a microbiome) to axenic or control Canton-S males. For these treatments, we measured fecundity across 54 h post-mating and transcript abundance at 6 h post-mating.

### 2.1 Both female and male microbiome status influence egg laying

Over the course of 54 h, control females mated to control males (CC) produced more eggs than axenic females, with an average of 77 eggs in CC crosses (± 4; sample size *n* = 50) versus 48 eggs in axenic females mated to axenic males (AA) (± 5; *n* = 49; *p* < 0.0001) and 53 eggs in axenic females mated to control males (AC) (± 6; *n* = 47; *p* = 0.0004) (Figure 1A). Interestingly, the CC cross produced significantly more eggs than the CA cross (control females mated to axenic males), which produced on average 51 eggs (± 4 *n* = 46; *p*= 0.003; Figure 1A). These results suggest that presence of a microbiome in the male has a positive impact on female fecundity.

**Figure 1.**
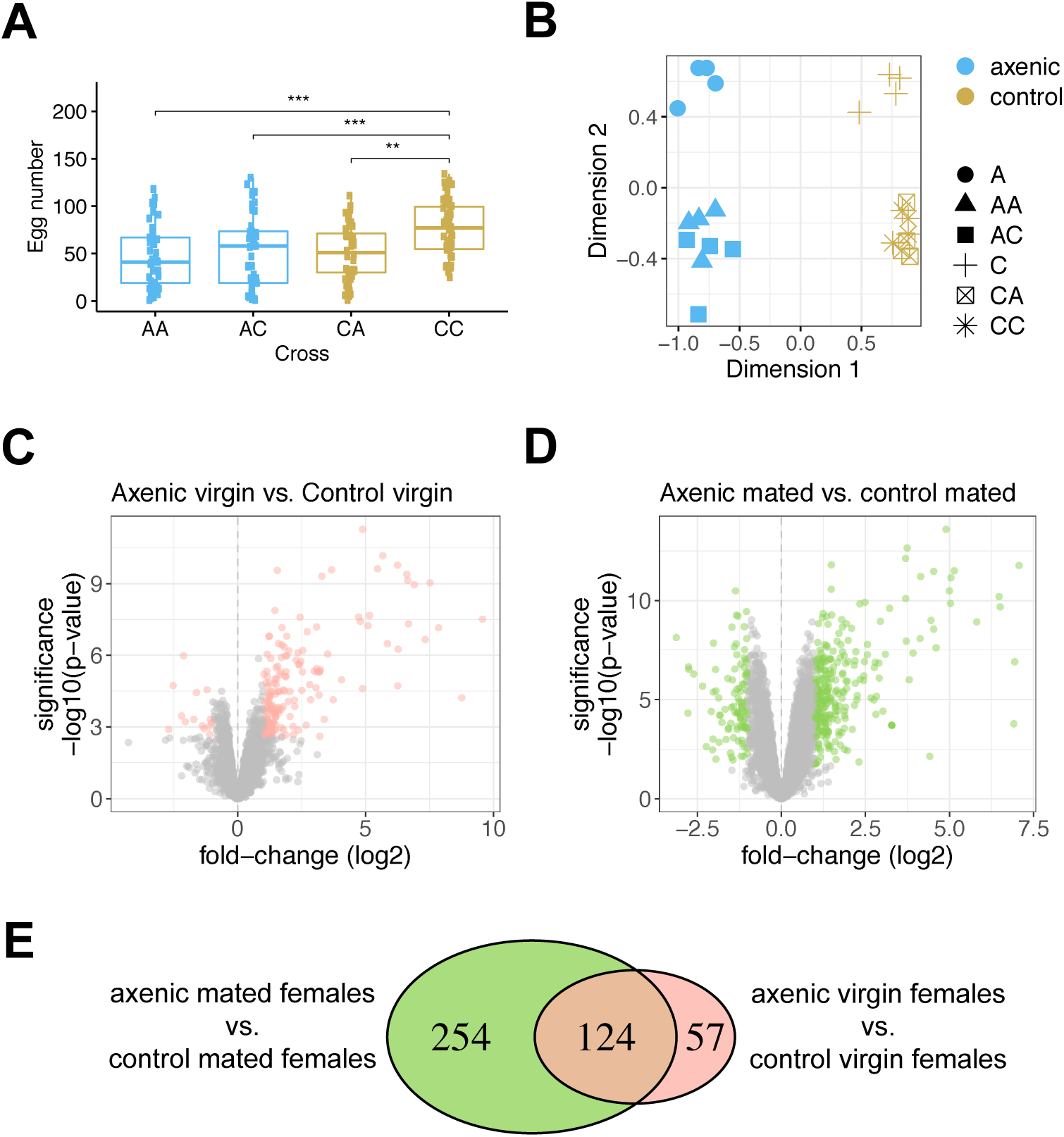
Fecundity and transcript abundance differences between axenic and control females. (A) Fecundity of axenic and control females after a single mating to axenic or control males. Each point represents eggs laid by one female over the course of 54 h. The first letter refers to the female’s microbiome status, the second letter refers to the male’s microbiome status (A=axenic, C=Control). Sample sizes: *n* = 49 for AA, *n* = 47 for AC, *n* = 46 for CA and *n* = 50 for CC. Groups were compared using a generalized linear mixed model with a Poisson response distribution. *** = p < 0.001; ** = p < 0.01. (B) Multidimensional scaling plot of replicates and samples used in the study. Microbiome status is represented by color, and sample origin represented by shape. (C) Volcano plot showing the results of a differential expression analysis of control virgin females relative to axenic virgin females. Significant genes (FDR < 0.05, >2-fold) are shown in pink. (D) Volcano plot showing the results of a differential expression analysis of control mated females relative to axenic mated females, averaged across the two male microbiome states. Significant genes (FDR < 0.05, >2-fold) are shown in green (E) Overlap of genes that are influenced by the microbiome in virgin and mated females.

### 2.2 Many transcripts differ in abundance between axenic and control virgin females, and many more are altered after mating

We investigated whether microbiome state influences a female’s transcriptome by directly comparing the transcriptomes of axenic and control females. First, we examined sample clustering using multidimensional scaling and found that axenic and control samples are clearly separated across the first dimension, while virgin and mated samples are separated across the second dimension (Figure 1B). Next, we compared transcript abundance between axenic and control virgin females and found 181 transcripts that differ in abundance (167 up-regulated and 14 down-regulated) (Figure 1C,E, Table S1). Finally, we compared transcript abundance between axenic and control mated females, irrespective of their mate’s microbiome status, and identified 378 transcripts that are differentially abundant (277 up-regulated and 101 down-regulated)(Figure 1D,E, Table S1). These two contrasts have 124 genes in common, suggesting that these genes constitute a “core” set that is influenced by the microbiome regardless of mating status (Figure 1E). In addition, 57 transcripts (51 up-regulated and 6 down-regulated) were affected by the microbiome specifically in virgin females, while 254 (161 up-regulated and 93 down-regulated) were affected by the microbiome specifically in mated females (Figure 1E).

### 2.3 The “core” set of microbiome response genes in females are involved in metabolic and immune processes and have an expression bias in the midgut

Among the 124 transcripts that are influenced by the microbiome state regardless of the females’ mating status, we found significant enrichment of Biological Process Gene Ontology (GO) terms related to the immune response and carbohydrate, nucleoside, lipid and amino acid metabolism (Figure S4A). Similarly, enriched Molecular Function GO terms included hydrolase, glucosidase, peptidase and lipase and sterol binding activity (Figure S4B). The majority of these transcripts (116/124) were up-regulated in control females relative to axenic females. These observations are in accordance with studies that showed that axenic flies have altered levels of glucose, trehalose, triglycerides, proteins and insulin-like signaling [30, 31, 53–56]. The core set of microbiome-responsive genes also have a clear expression bias in the female midgut (Figure 2A). This result is similar to that of [23], who found major transcriptome changes in the gut of axenic females, but detected few changes in non-gut tissues. Only eight transcripts had higher abundance in axenic females relative to control females. Of these, *Pka-R1* is noteworthy because it regulates feeding behavior [57] and PKA signaling acts downstream of dopamine signals to promote ovarian dormancy [58].

**Figure 2.**
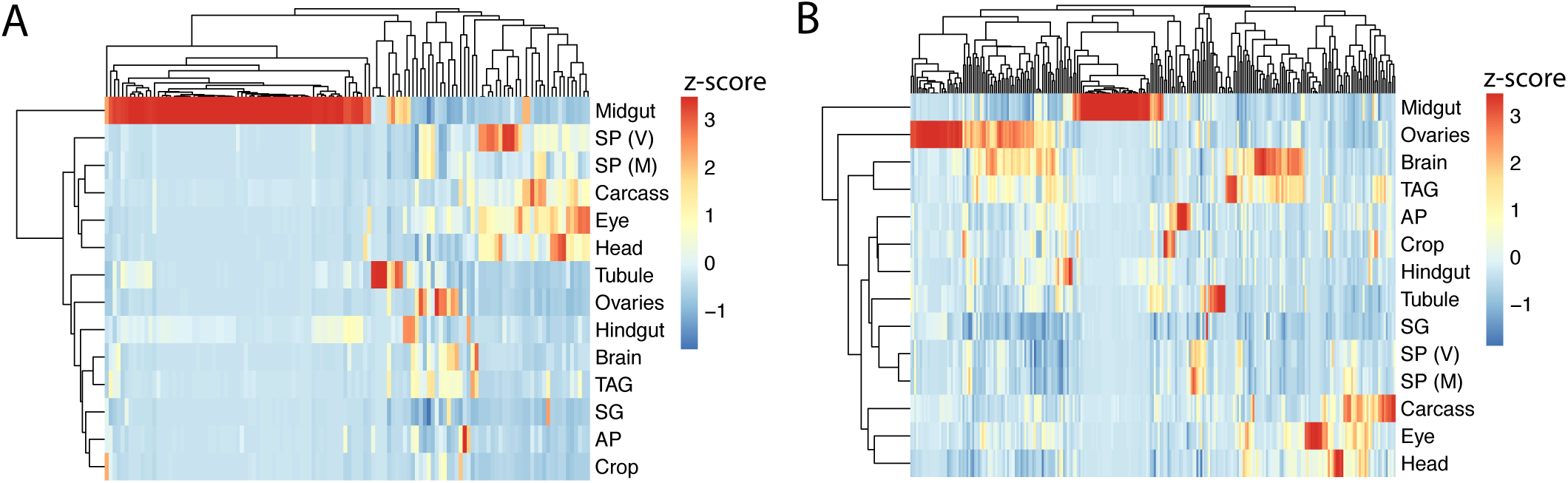
Tissue enrichment scores for differentially abundant transcripts across 14 female tissues. (A) Heatmap for 124 “core” genes whose RNA levels are influenced by the microbiome in both mated and virgin females. (B) Heatmap for 254 genes whose RNA levels are influenced by the microbiome in mated females only. Enrichment scores were calculated using TPM values from FlyAtlas2 [52]. (AP = anal plate; M = mated; SG = salivary gland; SP = spermathecae; TAG = Thoracico-abdominal ganglion; V = virgin)

In virgin females specifically, only 57 transcripts were affected by microbiome status (Figure 1C,E). Most of these (51/57) were up-regulated in control virgin females relative to axenic virgin females. These up-regulated transcripts were significantly enriched for peptidases and carbohydrate transmembrane transporters (Figure S4C). In addition, we identified 10 genes involved in the immune response and two genes involved in reproduction (*tj, Cp36*). Only six transcripts were down-regulated in control relative to axenic virgin females. One of these is *takeout*, which is associated with circadian rhythm, starvation and food intake [59]. We tested the transcript levels of *to* using qRT-PCR on independently collected samples and verified that it was downregulated in control virgin females relative to axenic virgin females (*p* = 0.005, Figure S6). We also performed a qRT-PCR on *Mtk* (an antimicrobial peptide) and *Tobi* (“target of brain insulin”; [60]). For both genes, we were able to validate a significant upregulation in control mated females relative to axenic mated females (Figure S6; *Mtk* p=0.04; *Tobi* p=0.007). *Mtk* and *Tobi* expression was not significantly higher in control virgin females relative to axenic virgin females in our qRT-PCR results.

### 2.4 Many of the 124 “core” genes influenced by the microbiome in this study were influenced by the microbiome in previously published studies

The study design we employed is different from the designs used by most published microbiome studies. Specifically, we created “control” flies with a conventional microbiome by adding homogenate of untreated flies onto the fly media, while most studies generate gnotobiotic flies which carry a limited, curated set of bacterial species that are usually found in the fly gut [21]. We created control flies to assess the effects of presence/absence of the microbiome, rather than the effects of specific bacteria. In addition, we did not want to omit potential effects of bacteria present in the reproductive tract, which have been shown to influence reproduction in several other species [61], but have not been characterized in *D. melanogaster*.

Despite the differences in study design, more than half (52%) of our “core” genes had been reported to be influenced by the microbiome in at least one of three other studies [22, 23, 25] (Table S2). Moreover, these 124 genes were enriched for similar functions, i.e., immune response and metabolic processes [22–25]. [23] also defined a “core” set of 152 genes whose transcript abundance was influenced by the microbiome in the female gut in flies with distinct genotypes (Oregon-R and Canton-S). We found only 11 genes that overlap between our core set and the core set from [23]. This likely reflects the use of gut versus whole fly, or it could be caused by differences in experimental design, such as growth conditions or fly genotypes. Still, these 11 genes fall into several broad functional classes that are affected by the microbiome both in this study and in [23], including immune and stress response genes (*AttA, AttB, GstD8*), genes affecting gut structure (*Mur29B, CG7017*), metabolism (*Npc2e, Acbp6, Gba1a, CG17192*) and gene expression (*CG15533*).

### 2.5 Transcripts involved in reproduction and neuronal function differ in abundance between axenic and control mated females

Next we analyzed the 254 transcripts that differ in abundance specifically between mated axenic and mated control females (Figure 1D-E). These transcripts have an expression bias to the midgut and the ovary, and to a lesser extent the brain (Figure 2B). Of the 254 transcripts, 161 transcripts were up-regulated and 93 transcripts were down-regulated in mated control females relative to mated axenic females (Figure 1D). We did not identify significantly enriched GO terms among the 161 up-regulated genes, but we detected genes associated with reproduction (29 genes; related GO terms included cell cycle, chromosome segregation, regulation of cell proliferation, stem cell population maintenance and macromolecule localization). We performed a Gene Set Enrichment Analysis (GSEA) using the fold-changes derived from the contrast between mated axenic females and mated control females, and identified multiple significant Biological Process terms associated with immunity, reproduction and metabolism (Table S3). Notably, for Molecular Function, only hydrolase activity was enriched (Figure 3A). Genes encoding hydrolases show a strong expression bias in the midgut and are largely composed of maltases and mannosidases (Figure 3B). Additional up-regulated genes include those involved in the immune response (*Def, PGRP-SC1b, PGRP-SD*), dopaminergic neurotransmission, (*Fer2, Catsup, Bx, Atpalpha*)(Figure 3C), and pigment biosynthesis (*yellow-f, bw*). Changes in pigment biosynthesis have been described in axenic flies [62], and this could reflect a sub-optimal metabolism in the absence of bacteria [63].

**Figure 3.**
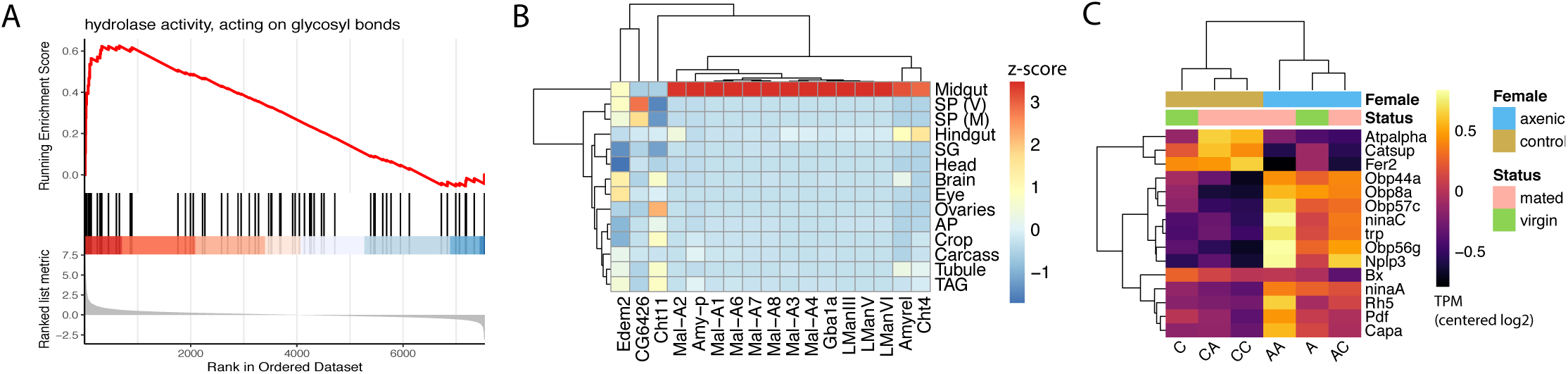
Genes whose mRNA levels are influenced by the microbiome in mated females have roles in metabolism and neuronal functions. (A) Gene Set Enrichment Analysis (GSEA) showing that transcripts involved in glycolitic metabolism are generally up-regulated in mated females with a microbiome relative to axenic mated females. The top panel of the GSEA plot shows the running enrichment score for a rank-orderd list of genes that are involved in hydrolase activity (genes are ranked based on the log2 fold change between mated control and mated axenic females, in decreasing order). Each vertical black line represents a gene that is involved in hydrolase activity. (B) Tissue enrichment Z-scores of genes encoding hydrolases, that are differentially expressed between mated axenic and mated control females. These genes show a strong expression bias in the female midgut (abbreviated tissue samples are the same as in Figure 2). (C) Heatmap showing mean centered log2 TPM for 14 genes with sensory or neuronal functions, whose transcript abundance is altered in mated females depending on whether they have a microbiome or not. (A = axenic, C = control). The first letter refers to the female’s microbiome status, the second letter refers to the male’s microbiome status. Female microbiome and mating status are indicated above the heatmap and in the key.

Among the 161 up-regulated genes, the ones with the highest up-regulation had at least a 6-fold higher transcript level in control compared to axenic mated females. These included five members of the histone H4 family (*His4:CG33891, His4:CG33893, His4:CG33895, His4:CG33897* and *His4:CG33899*). Histone gene expression occurs during S phase [64]. Thus, lower levels of histone transcripts suggest that fewer cells are dividing in axenic females compared to control females. This could reflect a reduction in oogenesis [37] or a reduction in cell proliferation in other tissues, such as the gut [23, 32].

The 93 transcripts with lower abundance in control mated females compared to axenic mated females were significantly enriched for genes involved in sensory perception (Figure S4D). These included four genes encoding odorant binding proteins (*Obp8a, Obp44a, Obp56g, Obp57c*), genes involved in phototransduction (*Rh5, ninaA* and *ninaC*) and a cation channel (*trp*) (Figure 3C). Furthermore, mRNAs encoding three neuropeptides (*Nplp3, Pdf, Capa*) and *TpnC4* and *TpnC41C*, which are part of the muscle troponin complex, were down-regulated in control mated females (Figure 3C).

### 2.6 22 genes respond to mating regardless of female microbiome status

We detected 22 transcripts that were up- or down-regulated in females after mating regardless of microbiome status (Table S4). We detected these transcripts by contrasting transcript abundance in mated females with that of the respective virgin females. Mating-responsive genes include three spermathecal serine-type endopeptidases (*Send2, CG17239, CG17234*); the metallopeptidase *Nep7*; *jhamt*, involved in juvenile hormone synthesis; a maltase, *Mal-B1*; a gene encoding an odorant binding protein *Obp83f*; *wbl*, involved in Toll signaling and dorso-ventral patterning; the antimicrobial peptide *Listericin*; and *CG14191*, which is involved in sarcomere function. Using qRT-PCR on independently collected samples, we confirmed the post-mating upregulation of *jhamt* in both axenic and control females (p-values for both contrasts < 0.001; Fig. S6). We further confirmed a downregulation of *Mal-B1* in control females after mating using qRT-PCR (p = 0.02; Fig. S6). *Mal-B1* transcript abundance was also lower after mating in axenic females, but that difference was not statistically significant based on the qRT-PCR data.

### 2.7 Male microbiome status does not affect post-mating transcript abundance in control females, but has a major effect on axenic females

Next we examined contrasts that reveal the effect of male microbiome status on the female’s post-mating transcriptome. When we directly compared transcript abundance in control females mated to control males with that of control females mated to axenic males, we did not detect any differentially-expressed transcripts (Figure 4A). This suggests that the male’s microbiome does not affect post-mating mRNA levels at six hours in females that have a microbiome. We then compared transcript abundance between axenic females mated to axenic males and axenic females mated to control males and found 136 transcripts that were differentially abundant (Figure 4A, Table S5). Hierarchical clustering of all samples based on the normalized expression for these 136 genes showed that the transcript abundance of these genes was similar across virgin females and mated control females; mated axenic females formed separate clusters depending on the male they mated with, showing opposite patterns of transcript abundance (Figure 4B). Only 14 transcripts were detected at a higher level in AC crosses relative to AA crosses. These included three immune effectors (*IM18, Dro* and *Listericin*; Figure 4C). The majority of the transcripts (122/136) were present at a lower level in AC crosses compared to AA crosses. These genes have an expression bias to ovaries and the midgut, brain and thoracico-abdominal ganglion (Figure S5). Gene Set Enrichment Analysis indicated a significant enrichment of peptidases among the 136 differentially expressed genes. Additional genes of interest identified using GO classification include 17 genes involved in the stress and immune response, 8 genes involved in reproduction (among which are *vas* and *jhamt*, Figure 4C) and genes with neuronal functions (*Atx2, NinaE, Bx, TBPH, Dsk*).

**Figure 4.**
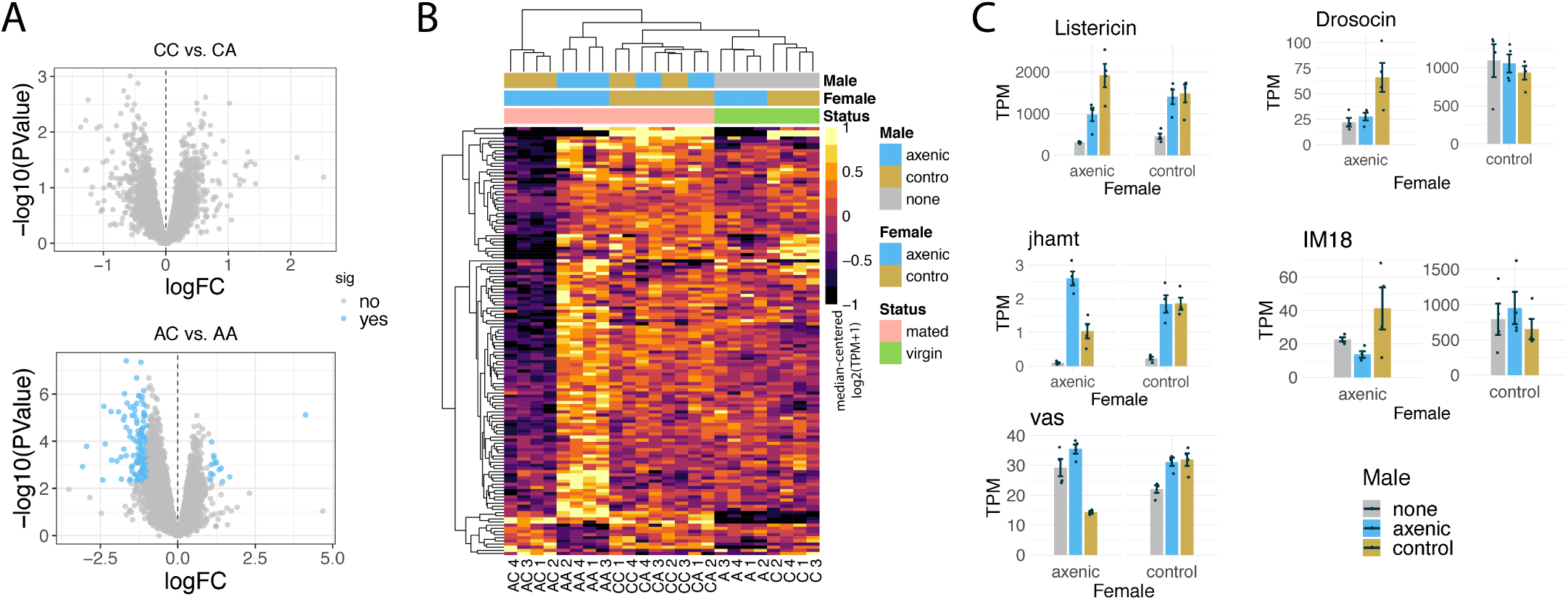
Analysis of female transcriptome changes that are influenced by the male’s microbiome status. (A) Volcano plot showing the results of a differential expression analysis assessing the effect of male microbiome status on control females (top) and axenic females (bottom). (B) Heatmap of normalized, batch-adjusted abundance values (TPM; transcripts per million) for transcripts that are altered in axenic females after mating with an axenic or control male. (C) Barplots of TPM values for a subset of genes that are influenced by the male’s microbiome in axenic mated females. Error bars represent standard error, and points represent triplicate TPM values.

## 3 Discussion

This study addressed two main questions: (1) Is a female’s transcriptome influenced by interactions between her microbiome and her mating status? and (2) can interactions between the female’s microbiome and the male’s microbiome influence the female’s transcriptome and fecundity? We found evidence for such interactions and discuss their implications for the female’s reproduction and metabolism, neuronal functions and immune gene expression.

Using a fecundity assay, we observed that females without a microbiome laid fewer eggs than females with a microbiome. This observation confirms published results [28, 37, 39]. Furthermore, we observed lower mRNA abundance of genes involved in egg production in axenic females. This was apparent in axenic virgin females, which had—compared to control virgin females—lower mRNA levels of *Cp36*, which encodes a chorion protein [65], and *tj*, which is involved in gonad morphogenesis [66]. After mating, we detected differential abundance of many additional transcripts involved in reproduction. This is likely because mating, and specifically seminal fluid proteins, kickstart egg production [1].

We observed an up-regulation of *jhamt* after mating in all females, whether axenic or control. JHAMT is essential for juvenile hormone (JH) synthesis [67]. JH is an endocrine factor that stimulates the production of vitellogenin and yolk proteins [68] and JH also suppresses the mated fly’s ablity to resist infection [69]. JH production is stimulated by the male seminal fluid protein Sex Peptide [70]. The observed up-regulation of *jhamt* mRNA suggests that, in the absence of a microbiome, signals received during mating still elicit an attempt to initiate oogenesis via JH, but somehow oogenesis is curtailed in the absence of a microbiome. One factor that likely contributes is an altered metabolism in axenic vs. control mated females, which was observed both in our study (using transcript abundance) and previously published studies (using transcript abundance or metabolite measurements) [22–26, 30, 31]. Our study took these observations further, by showing that the abundance of mRNAs of metabolic genes is lower in the absence of the microbiome both before and after mating. Studies of the post-mating responses in *D. melanogaster* females have shown that mating induces metabolic changes and changes in feeding behavior, likely to accommodate the high energy demands of oogenesis [19, 20, 71–74]. Thus, in the absence of a microbiome, a females’ ability to manage the metabolic changes needed to sustain egg production might be negatively affected.

Our data further show that the microbiome influences the mRNA abundance of genes with neuronal functions. This class of genes was also reported as influenced by the microbiome by [25]. Our data show that this phenomenon specifically occurred in mated females and not in virgin females. We observed an up-regulation of genes encoding odorant binding proteins and genes encoding components needed for phototransduction in axenic mated females relative to control mated females. Several studies have shown that fly olfactory behavior changes in the absence of a microbiome or upon changes in microbial composition [34, 39, 75, 76], but none reported effects on vision. Transcript abundance of genes involved in olfaction and phototransduction also change after mating in *Drosophila* [4, 5, 8, 14] and honeybees [77]. In flies, such changes in sensory genes after mating could mediate changes in female receptivity to other males [78, 79] or aid her in finding suitable sites for egg laying [18, 80]. Thus, it is possible that post-mating sensory responses are altered depending on female microbiome state.

We further observed changes in the mRNA levels of two genes with functions in circadian and locomotor behavior, Bx and Pdf [81, 82]. In addition, two troponins required for muscle contraction had higher mRNA levels in mated axenic females relative to mated control females. *D. melanogaster* female activity levels increase after mating [83] and in the absence of a microbiome [35]. Thus, the transcript changes we observed could reflect those changes in locomotion on a molecular level. In addition, we observed changes in the mRNA levels of genes involved in dopamine signaling (*Fer2, Catsup, Bx, Atpalpha*) [84, 85]. Dopamine has many effects on fly behavior [86] and the causes and consequences of changes in dopamine signaling cannot be determined based on the current study. However, these results indicate that interactions between the female’s microbiome and her mating status have a significant impact on mRNA levels of neuronal genes.

A common observation in studies of the female post-mating response is that immunity genes are up-regulated after mating [2, 4, 10, 87], but the basis for this post-mating induction of immune response genes is not fully understood [51]. Here we found that transcripts of immunity genes were up-regulated in control females relative to axenic females, confirming results from other studies [22–26]. In addition, the transcript levels of some immunity genes were influenced by interactions between the female’s microbiome and her mating status. For example, *Def, PGRP-SC1b* and *PGRP-SD* had higher mRNA levels in control mated females relative to axenic mated females, while this was not the case when comparing virgin females. This indicates that mating elevates the mRNA levels of these genes only in the presence of a microbiome in the female. On the other hand, transcripts of the antimicrobial peptide *Listericin* were up-regulated by mating in all females, regardless of whether they or their mate had a microbiome. Listericin expression has been reported to be regulated by PGRP-LE and JAK-STAT signaling [88]. Our observation is particularly interesting because it suggests that some aspect of mating, without the need for microbiota, can increase the RNA levels of this antimicrobial peptide, perhaps by activating JAK-STAT signaling rather than Toll and imd signaling, the canonical signaling pathways in response to septic threats. This aspect of mating could be copulatory wounding [89, 90], or exposure to sperm or seminal fluid proteins [4–7, 17, 18, 91]. If an axenic female mated to a control male, additional immune transcripts were up-regulated relative to when an axenic female mated to an axenic male (e.g. *IM18, Dro*, and *Listericin*), indicating that exposure to microbiota during courtship or copulation stimulates an additional up-regulation of these immune gene transcripts. There is increasing attention for the role of reproductive tract microbiota in reproductive success [61, 92], and a female- and male-specific reproductive tract microbiome has been characterized in *Anopheles* mosquitos, but whether *D. melanogaster* have reproductive-tract specific microbiomes that can influence the post-mating up-regulation of immune transcripts is not yet known.

Using RNA-seq data, we did not observe effects of male microbiome status on the transcriptome of control females, but we observed significant effects on the transcriptome of axenic females. We wondered whether exposure to bacteria on the male’s cuticle during courtship or mating, or exposure to male excreta, could make the mRNA levels of an axenic female more similar to those of a control female. However, at six hours after mating, that does not appear to be the case. The genes whose mRNA levels were influenced by the male’s microbiome had various functions (including egg production) and were mostly down-regulated after mating with a control male. Perhaps a sudden exposure to bacteria during mating does not make axenic female mRNA levels more similar to those of control females because resources are used to initiate an immune response rather than oogenesis. Additional experiments at multiple time points would be necessary to resolve this hypothesis.

Aside from transcript abundance, female fecundity was also influenced by the male’s microbiome. Fecundity was lower not only in axenic females, but also in control females that had mated with an axenic male. This indicates that the absence of a microbiome impacts a male’s reproductive success. Interactions between a male’s reproductive success and his microbiome were also observed by [38]. For example, Morimoto et al. (2017) observed that gnotobiotic males carrying only *Lactobacillus plantarum* had a longer copulation duration and induced higher short-term egg laying in their mates. Axenic males could differ from control males in pheromone production, or in the production, transfer or quality of seminal fluid proteins or sperm. The reduced fecundity in control females mated to axenic males was not accompanied by transcript level changes in our dataset, possibly due to the time point measured. It is also possible that egg production is unaffected in control females mated to axenic males, but that they differ from control females mated to control males in their frequency of egg deposition.

To conclude, we have shown that a *D. melanogaster* female’s transcriptome is influenced by interactions between her microbiome and her mating status, and that both transcript abundance and fecundity are influenced by interactions between the female’s microbiome and that of her mate. Our results demonstrate the importance of considering a females’ mating status to better understand and interpret the host microbiome’s impact on overall fitness.

## 4 Materials and Methods

### 4.1 Fly stocks, rearing and the generation of axenic and control flies

Canton-S flies were maintained at 25*°*C on yeast-sucrose-cornmeal food (7 g agar; 12 g yeast; 12 g cornmeal; 40 g sucrose; 1,000 ml water, 26.5 ml Tegosept; 12 ml acid mixture) in a 12 h light/dark cycle. To generate axenic and control files, we followed the protocol described by [93]. Briefly, population cages of Canton-S flies were set up and females were allowed to oviposit on grape juice agar plates for 2-3 days until robust egg-laying began. On the third day, embryos were collected and treated twice (2.5 min each time) with a 0.6% sodium hypochlorite solution and triple-rinsed in autoclaved distilled water in a biosafety cabinet. Axenic embryos were allowed to hatch in 50 ml sterile vials containing yeasted autoclaved food with 40 ul of 1X PBS added to the food surface. To generate controls, the same embryo dechorionation procedure was followed, but the tubes for the control samples received 40 ul of Canton-S fly homogenate (prepared in aliquots of 200 ul, at a concentration of 50 flies in 200 ul 1X PBS) on the food surface to add the full set of bacteria found in our lab’s Canton-S stock. After seven days, pupae from axenic and control tubes were collected and twice treated with 0.6% sodium hypochlorite for 30 sec and subsequently rinsed three times in autoclaved distilled water. Pupae were individually placed into a vial with sterile food. Each vial with a 7-day old control pupa received 20 ul of Canton-S fly homogenate on the food surface (on average 2.5 flies ground up for each vial). Each vial with an axenic pupa received 20 ul of sterile 1X PBS.

### 4.2 Confirmation of microbiome status

We performed two assays to ensure axenic flies were germ-free and to ensure the presence of a microbiome in control flies: 1) Individual flies were homogenized in De Man, Rogosa and Sharpe (MRS) broth and plated on MRS plates as in [93], which were then incubated at 29*°* C for 2-3 days and checked for colonies, and 2) a PCR assay was performed according to the methods in [30]. Briefly, genomic DNA was extracted from 3-10 pooled axenic or control larvae, pupae or adult flies. PCR was run using primers designed by [30] for a conserved region of the bacterial 16S rRNA gene. PCR products were run on a 1% agarose gel to confirm the absence of bacteria in axenic individuals and the presence of bacteria in controls (Figure S1). We also compared levels of bacteria in 9-12 pooled adults of our control flies with levels of bacteria in 9-12 pooled untreated (not dechorionated) adult Canton-S flies and found an enrichment of bacteria in our control flies (Figure S1). Absence of *Wolbachia pipientis* in our lab’s Canton-S stock was confirmed using a PCR assay described by [94].

### 4.3 Mating assay and sample collection for RNA-seq and qRT-PCR

Five day old virgin flies were used for the mating assays. Axenic and control females and males were singly mated in a 2×2 full factorial design: control females x control males (CC), control females x axenic males (CA), axenic females x axenic males (AA) and axenic females x control males (AC). Matings were observed and males were removed from the vial after copulation ended. The end time of copulation was recorded and females were flash-frozen six hours after mating, at which time virgin axenic and control females from the same cohort were also flash-frozen. For RNA-seq, we collected four replicates for each of the six treatments on the same day. Around ten females were pooled per replicate. To carry out qRT-PCR confirmations using samples independent from those used for the RNA-seq, we used flies produced from eggs that were dechorionated on a different day from the ones used for the RNA-seq samples. Females were mated to males of the same microbiome status as themselves and were flash frozen six hours after mating, at which time virgin females were also frozen. For each treatment for the qRT-PCR, three replicates of 10 pooled females were collected.

### 4.4 RNA extraction, RNA-seq library preparation and qRT-PCR methods

To extract whole RNA, a pool of ∼10 frozen females from each sample was homogenized in TRIzol following manufacturer’s guidelines (Thermo Fisher Scientific Inc., MA). Following liquid phase separation, the RNA-containing upper layer was subjected to column purification and DNase treatment using the RNeasy Mini Kit (Qiagen inc., MD). Purified RNA was quantified and saved at −80*°* C for library preparation. RNA-seq libraries were made using the Lexogen 3’ FWD kit following the manufacturer’s protocol (Lexogen, NH). Libraries were quantified on an Agilent 2100 Bioanalyzer before pooling and cluster generation/sequencing on an Illumina NextSeq platform.

For qRT-PCR, RNA was extracted as above. RNA was DNase treated using RQ1 RNase-Free DNase (Promega, WI) and cDNA was synthesized using SMARTScribeTM Reverse Transcriptase (Clontech, CA). qRT-PCR reactions were run on three biological replicates, each with three technical replicates, on a Roche LightCylcer 480 Instrument II using LightCycler 480 SYBR GreenI Master (Roche, NJ). Primers were designed using Primer Blast, except for the gene *jhamt*, for which we used primers designed by [69] and we verified that primer efficiency was above 80%. Primer sequences can be found in Table S6. *Rp49* or *Nervana* were used as control genes. We verified that these genes were not among the differentially expressed genes for the contrasts of interest. Ct values were analyzed using linear models in R. For each of the five genes tested, we set up an independent linear model using Ct value of the gene of interest as response variable, and using “sample” (A, C, AA or CC) and Ct value of the resp. housekeeping gene as explanatory variables (both fixed effects). The linear models were run on three biological replicates for each gene tested. Each biological replicate was the average of three technical replicates. After fitting the models, we calculated estimated marginal means (EMM), standard error of the EMM and pairwise contrasts between samples using the R package emmeans (Searle et al. 1980). P-values of pairwise contrasts were corrected for multiple testing using the Benjamini-Hochberg method (Benjamini and Hochberg 1997). We used a Shapiro test to ensure that residuals of the fitted models followed a normal distribution and used a Levene’s test to ensure homogeneity of variance of the Ct values.

### 4.5 Read processing, alignment and differential expression analysis

Raw reads were processed by trimming 10 bases from the 5’ end and quality trimming from the 3’ end to a minimum quality PHRED score of 20. Processed reads were mapped to the *D. melanogaster* transcriptome (Flybase r6.23) with bowtie2, and read counts and normalized abundances were extracted using eXpress [95, 96]. All differential expression analyses were performed in R using the packages EdgeR [97] and RUVseq [98]. We filtered genes with cpm <1 in at least 4 samples, leaving 7,649 genes in the dataset. After normalizing counts based on library size, a clear batch effect was visible (Figure S2A). We used RUVseq to identify *k*=3 additional variables that were added to the linear model in EdgeR. These variables were estimated by RUVseq based on the residuals from a linear model fitted with the sample variables. Adjusting for three additional unknown variables resulted in improved clustering of samples in a PCA plot (Figure S2B) and improved Pearsons’s correlations between replicates of the same sample.

We set up contrasts to 1) identify changes in transcript abundance in the female that depend on her microbiome state, 2) identify mating-responsive transcripts in females, and 3) identify changes in transcript abundance in females that are influenced by the microbiome state of her mate. Transcripts were considered significantly differentially abundant if the change was >2-fold and had a p-value adjusted for multiple testing <0.05 [99]. The package ClusterProfiler [100] was used for Gene Ontology (GO) and Gene Set Enrichment Analysis (GSEA). GSEA was based on fold changes for all 7,649 genes in the filtered dataset, using a minimal gene set of 50 genes and a maximal gene set of 500 genes. GO enrichment analysis was performed on the genes with differentially abundant transcripts, using a minimal gene set of 2 genes, and using all 7,649 genes as background.We called a GO category as significantly enriched if it had an adjusted p-value ≤0.05. DAVID [101, 102] and Flybase [103] were queried for further functional annotation of genes.

### 4.6 Tissue enrichment calculation

To determine if differentially abundant transcripts had an expression bias to particular female tissues, we used a custom analysis of gene expression data from the FlyAtlas (version 2) database [52] (https://github.com/YazBraimah/FlyAtlas2). We calculated tissue enrichment by dividing the normalized expression value (in transcripts per million, or TPM) of the gene of interest in the tissue of interest by the TPM value for that gene in the whole female body. As described on http://flyatlas.gla.ac.uk/FlyAtlas2/index.html?page=help#, when whole body TPM values were <2, we set them to 2 for the enrichment calculation.

### 4.7 Fertility assay

We performed a fertility assay by measuring the number of eggs produced by axenic (A) and control (C) females that were mated to axenic or control males. Matings for the egg laying assay were performed as described above, with sample sizes (female designated first in the cross): *n* = 49 for AA, *n* = 47 for AC, *n* = 46 for CA and *n* = 50 for CC. These sample sizes exclude females that did not survive or escaped during the assay. At the end of copulation, which occurred in vial 0 (V0), males were removed and females were transferred into a new vial (V1), in which they were allowed to lay eggs for six hours. After six hours, females were moved to V2 for 24 h, then transferred to V3 for 24 h, after which the females were discarded. Each time females were transferred to a new vial, egg number in the previous vial was recorded. Fly food in V0 was prepared as described above for the RNA-seq assay. V1, V2 and V3 contained the same autoclaved food as described above, but without the addition of yeast or fly homogenate. For each time point, we assessed the presence or absence of bacteria in 2-3 pooled flies using PCR for bacterial 16S rRNA. At each time point, bacteria were absent in axenic flies. Control flies contained bacteria at each time point, but the amount decreased with each transfer onto sterile food that did not contain fly homogenate (Figure S3). The total number of eggs produced by each female was analyzed in R using a generalized linear mixed model with an assumed Poisson response distribution (lme4; [104]), with fixed effects for female microbiome status, male microbiome status and their interaction, an observation-level random effect to account for overdispersion [105] and a random effect to account for the person counting the eggs. The package emmeans (https://cran.r-project.org/web/packages/emmeans/index.html) was used to calculate *p*-values for pairwise comparisons between the four treatments (corrected for multiple testing). Count data are available in Table S7.

### 4.8 Data and script availability

The raw Illumina short-read sequences are available through the Sequence Read Archive (SRA) under project accession PRJNA629997. The analysis scripts are available as a GitHub repository (https://github.com/YazBraimah/Axenic.PM).

## Acknowledgments

We thank the Feschotte and Johnson labs at Cornell for sharing equipment, E. Cosgrove for assistance with RNA-seq batch effects, J. Bubnell for help with *Wolbachia* detection, A. Douglas and members of the Clark and Wolfner labs for comments and discussion, and NIH RO1 HD059060 to MFW and AGC for funding.

## Author contributions statement

SYND, YHA-B, MFW and AGC conceived the experiments. SYND and YHA-B conducted the experiments. SYND and YHA-B analysed the results. All authors reviewed the manuscript.

## Supporting Information

**Figure S1.**
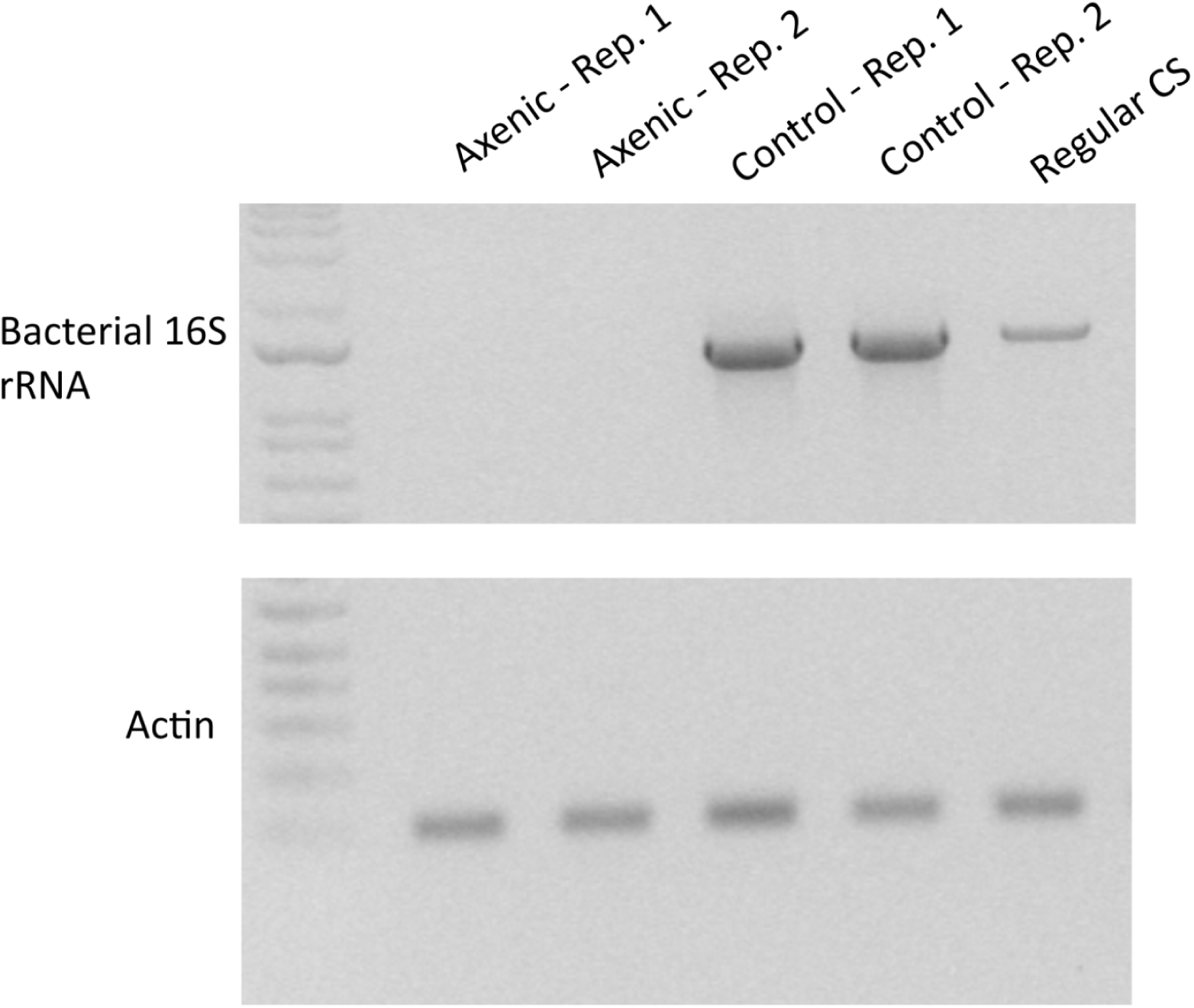
PCR assay to verify absence of bacteria in axenic flies and presence of bacteria in control. A mix of 9-12 pooled adult females and males was used for each sample on this gel. Axenic = dechorionated Canton-S flies with sterile 1X PBS added to sterile food. Control = dechorionated Canton-S flies with fly homogenate added to sterile fly food. Regular = untreated Canton-S flies. Rep. = biological replicate.

**Figure S2.**
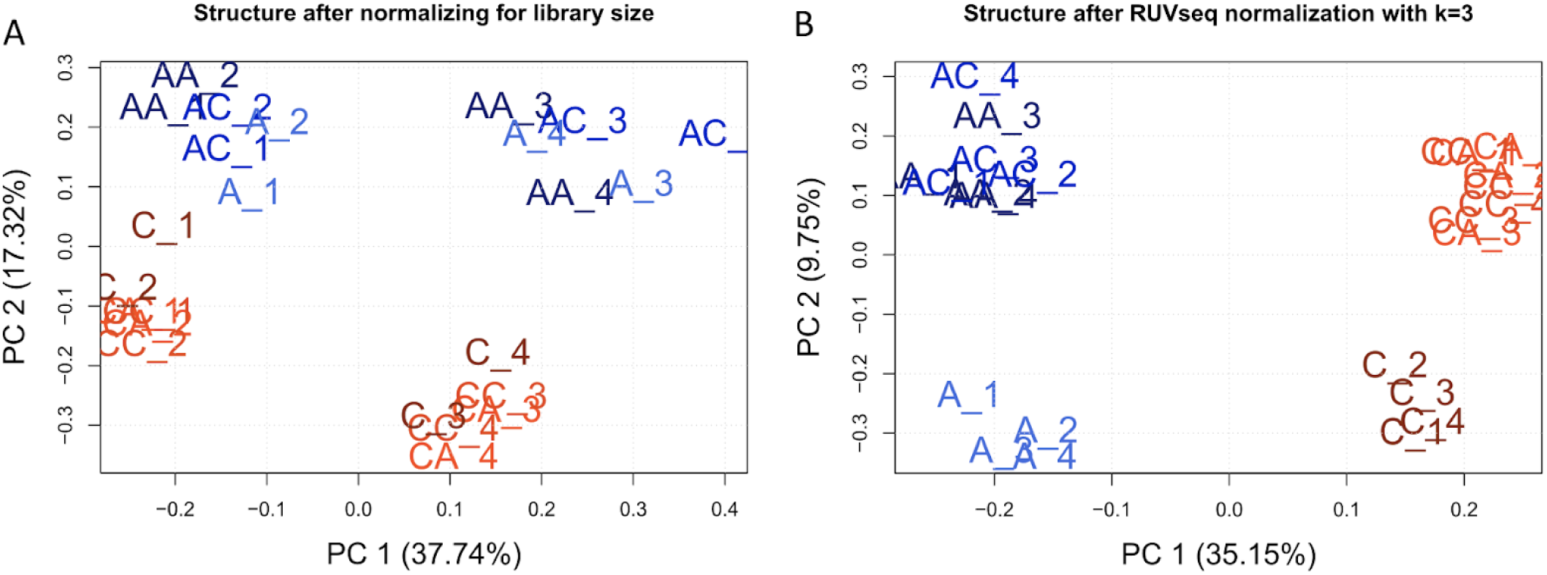
Principal Component Analysis of RNA-seq samples. A: Before adjustment for batch effect using RUVseq. B: After adjusting for batch effect using RUVseq. AA = axenic female mated to axenic male; AC = axenic female mated to control male; CA = control female mated to axenic male; CC = control female mated to control male, C = control virgin female, A = axenic virgin female. Numbers indicate the batch (biological replicate).

**Figure S3.**
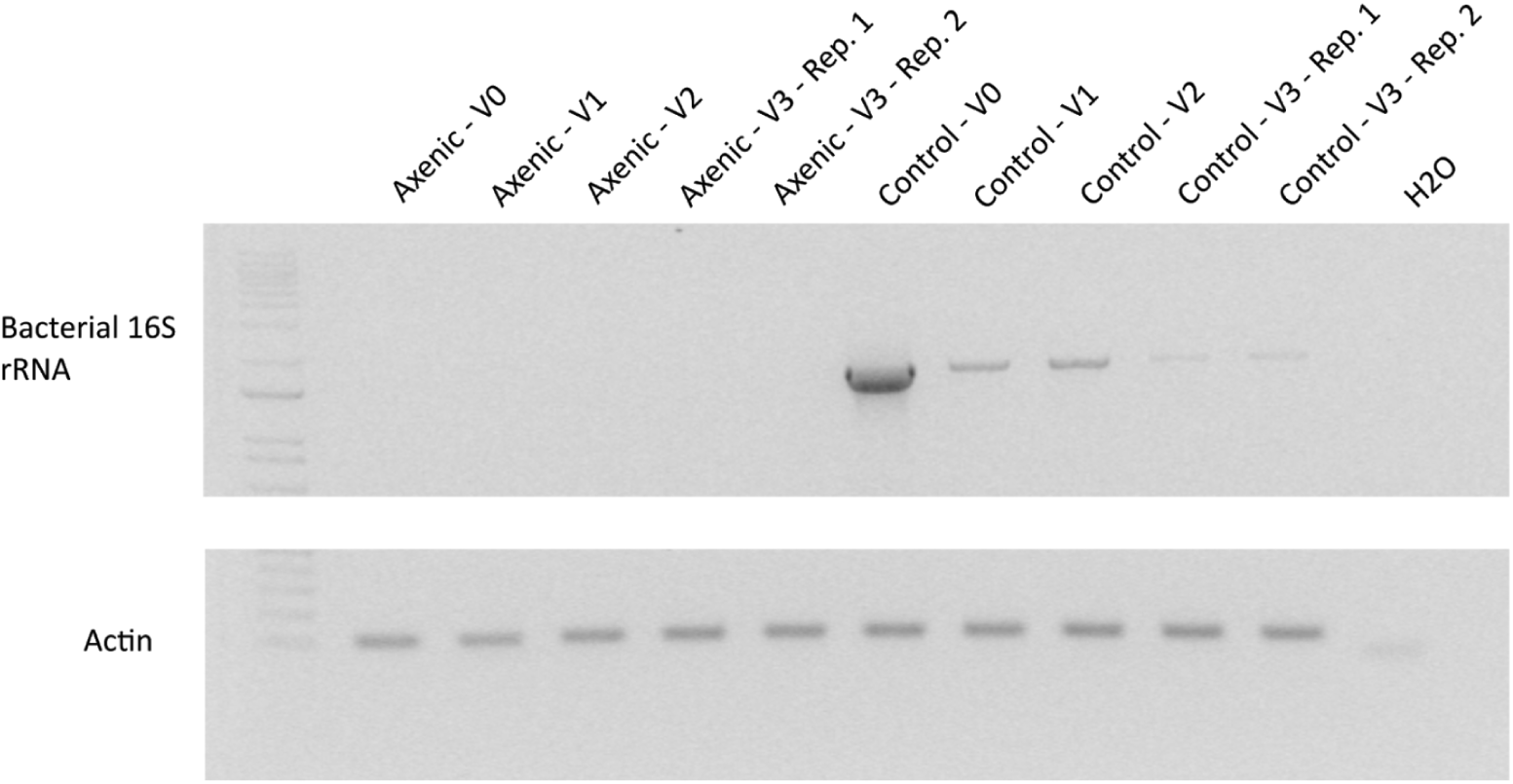
PCR assay to verify the presence of bacteria in control flies and absence of bacteria in axenic flies used for the egg laying assay. 2-3 flies were pooled at each time point. V0-3 = vial 0 to 3. Rep. = biological replicate.

**Figure S4.**
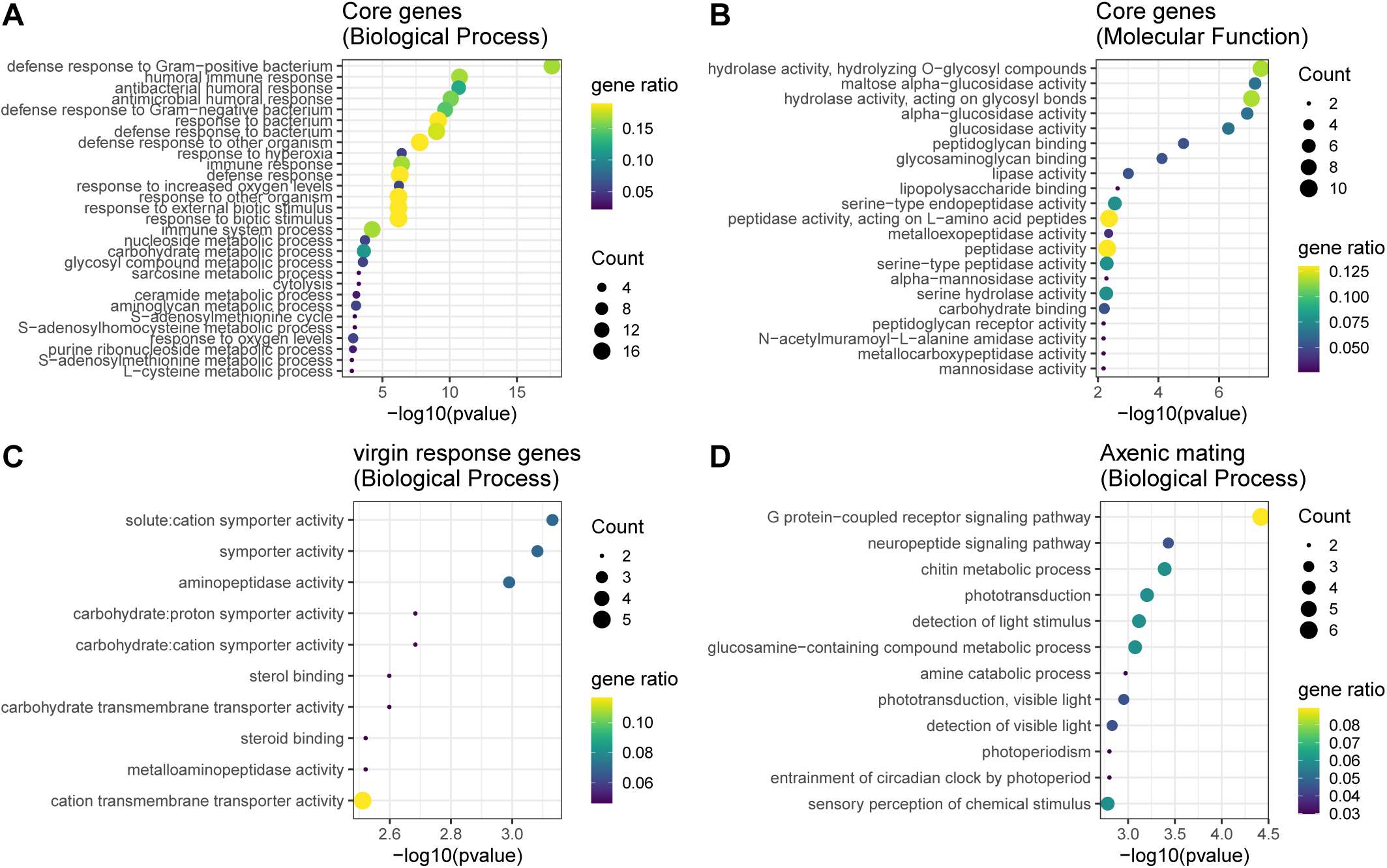
GO term enrichment analysis for differentially abundant transcripts. (A-B) Enriched GO terms for the 124 “core” genes whose RNA levels are influenced by the microbiome in both virgin and mated females. (C) Enriched Biological Process GO terms for 57 genes whose RNA levels are influenced by the microbiome in virgin females only. (D) Enriched Biological Process GO terms for 93 genes that are down-regulated in mated axenic females relative to mated control females.

**Figure S5.**
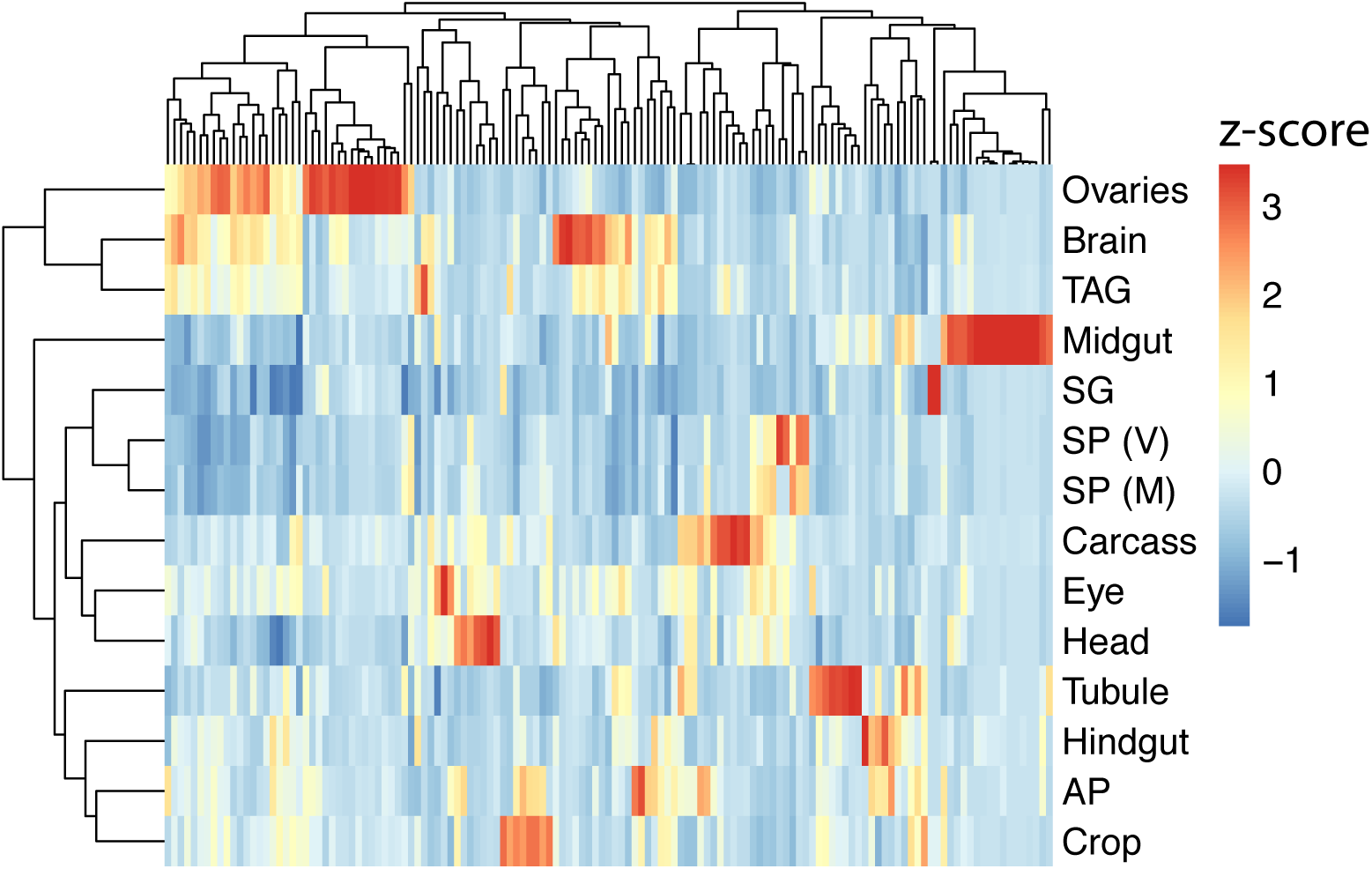
Enrichment values across 14 female tissues for genes whose transcript abundance is impacted in axenic females depending on the male’s microbiome. Enrichment scores were calculated using TPM values from FlyAtlas2 [52]. (AP = Anal plate; SP = spermathecae; M = mated; V = virgin; TAG = thoracico-abdominal ganglion)

**Figure S6.**
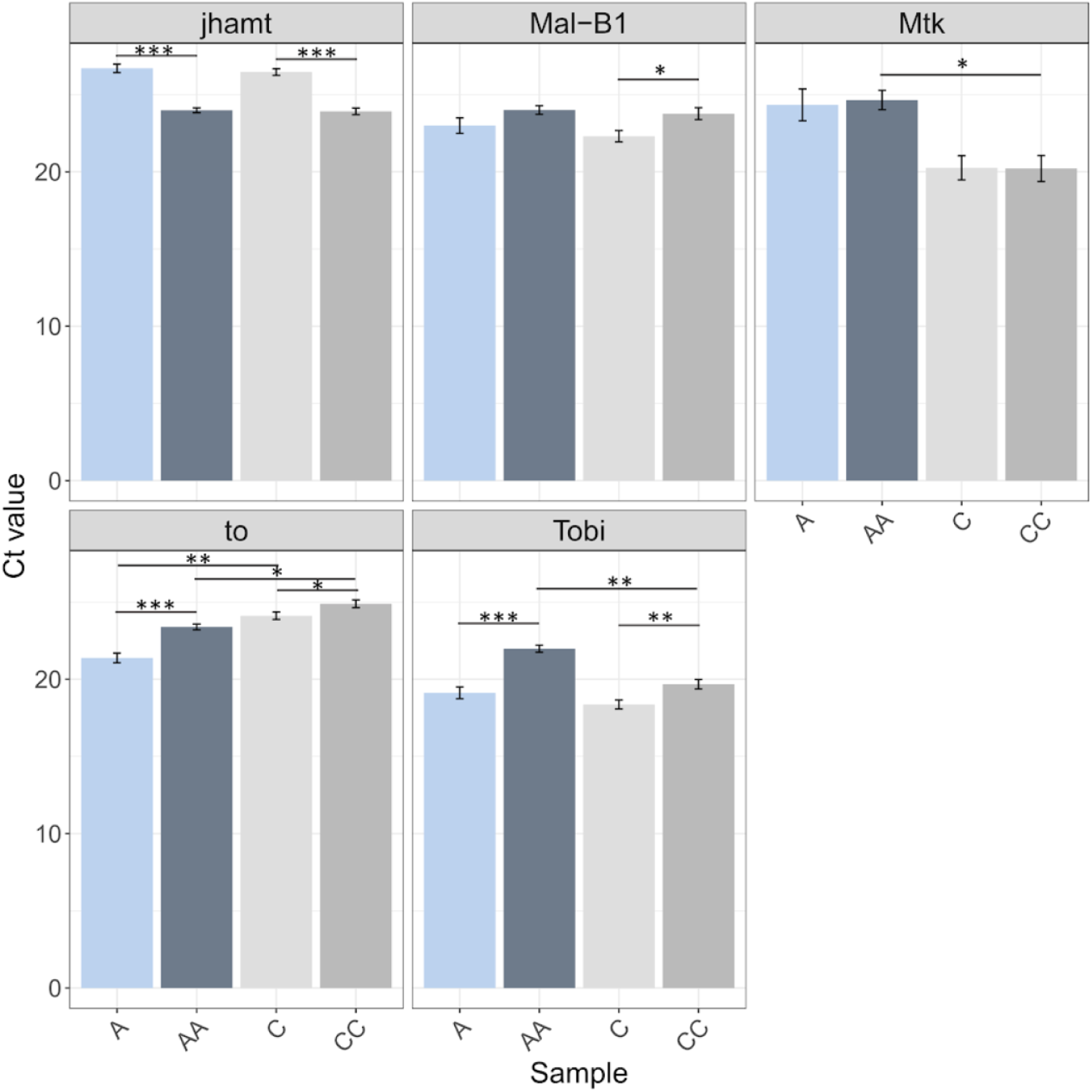
qRT-PCR validation of changes in transcript abundance observed in the RNA-seq analysis. Samples for qRT-PCR were collected independently from those used for RNA-seq. A = axenic virgin female; C = control virgin female; AA = axenic female mated to axenic male; CC = control female mated to control male. Bars show the estimated marginal means (EMM) of Ct values and standard errors of the EMM, based on linear models that were fitted for each gene independently. A higher Ct value indicates a lower expression of the gene. Each EMM was determined based on 3 biological replicates, which each contained 10 pooled females. All expression values were normalized to the housekeeping gene *Nervana*, except for *Mal-B1*, which was normalized against *Rp49*. Stars indicate significance of pairwise contrasts between the samples. * p < 0.05; ** p < 0.01; *** p < 0.001. If no star is shown, the contrast was not significant. After correcting for multiple testing using the Benjamini-Hochberg correction, no contrasts are significantly different for *Mtk*; for *jhamt*, both A-AA and C-CC have *p* < 0.05; for *to*, A-AA and A-C have *p* < 0.05; for *Tobi*, A-AA, AA-CC and C-CC have *p* < 0.05 and for *Mal-B1*, C-CC has *p* < 0.05.

